# Enzyme capacity-based genome scale modelling of CHO cells

**DOI:** 10.1101/2019.12.19.883652

**Authors:** Hock Chuan Yeo, Jongkwang Hong, Meiyappan Lakshmanan, Dong-Yup Lee

## Abstract

Chinese hamster ovary (CHO) cells are most prevalently used for producing recombinant therapeutics in biomanufacturing. Recently, more rational and systems approaches have been increasingly exploited to identify key metabolic bottlenecks and engineering targets for cell line engineering and process development based on the CHO genome-scale metabolic model which mechanistically characterizes cell culture behaviours. However, it is still challenging to quantify plausible intracellular fluxes and discern metabolic pathway usages considering various clonal traits and bioprocessing conditions. Thus, we newly incorporated enzyme kinetic information into the updated CHO genome-scale model (*i*CHO2291) and added enzyme capacity constraints within the flux balance analysis framework (ecFBA) to significantly reduce the flux variability in biologically meaningful manner, as such improving the accuracy of intracellular flux prediction. Interestingly, ecFBA could capture the overflow metabolism under the glucose excess condition where the usage of oxidative phosphorylation is limited by the enzyme capacity. In addition, its applicability was successfully demonstrated via a case study where the clone- and media-specific lactate metabolism was deciphered, suggesting that the lactate-pyruvate cycling could be beneficial for CHO cells to efficiently utilize the mitochondrial redox capacity. In summary, *i*CHO2296 with ecFBA can be used to confidently elucidate cell cultures and effectively identify key engineering targets, thus guiding bioprocess optimization and cell engineering efforts as a part of digital twin model for advanced biomanufacturing in future.

## 1. Introduction

Chinese hamster Ovary (CHO) cells are the most prevalent mammalian hosts for manufacturing therapeutic proteins, accounting for more than 80% of all monoclonal antibodies in the market (Walsh, 2018). Such widespread use of CHO cells is mainly due to the capability for post-translational modifications (e.g. *N-glycosylation*), ease of genetic manipulation, and the potential to thrive in suspension cultures. Although the production titre of CHO cells has increased by up to a thousand times over the decades, their efficiency can be further enhanced at two different levels: 1) cell line development and engineering, and 2) bioprocess development by identifying critical process parameters (CPPs), designing cell culture media and feeding strategies for the enhanced key performance indicators (KPIs), e.g., titre and yield, of the cell culture (Kyriakopoulos et al., 2018; Hong et al., 2018a). However, industrially most of such advances to date have been achieved only through the application of empirical techniques, such as ‘design of experiments’ (DOE) method, and black-box statistical approaches which are mainly based on historical bioprocess data sets; similarly, a priori knowledge- or experience-based cell line development and engineering are also being undertaken in ad-hoc manner. Thus, it is now highly required to develop more rational approaches to better understand CHO cell physiology, thereby enhancing their KPIs (Kildegaard et al., 2013). In this regard, the recent availability of CHO cell line-specific genome sequence information facilitated the development of systematic framework whereby *in silico* mechanistic models can be combined with increasingly available multi-omics datasets in order to describe cell culture behaviours. Previously, a genome scale metabolic model (GEM) of mouse (Selvarasu et al., 2010) was adopted to identify limiting nutrients in the growth of CHO fed-batch culture (Selvarasu et al., 2012), and to compare metabolic differences associated with the switching of lactate production to uptake in CHO cells (Martínez et al., 2013). Then, the first-ever CHO GEM was reconstructed through world-wide community efforts (Hefzi et al., 2016). The availability of such CHO GEM has greatly aided the characterization of various CHO cell line specific traits (Hefzi et al., 2016; Lakshmanan et al., 2019; Pan et al., 2017; Yusufi et al., 2017) and several bioprocessing conditions (Calmels et al., 2019; Hong et al., 2020; Park et al., 2018; Vodopivec et al., 2019).

Various *in silico* approaches have been employed to describe CHO cell metabolism, which include kinetic modelling, metabolic flux analysis (MFA) and constraint-based flux balance analysis (FBA) (Galleguillos et al., 2017; Huang et al., 2017; Kyriakopoulos et al., 2018). Of them, constraint-based FBA has been widely applied for analysing cellular metabolism at genome-scale as it is amenable to large-scale model, requiring only reaction stoichiometry and the mass balance of metabolites. While the utility of FBA has been successfully demonstrated by understanding cellular behaviour of CHO cells via aforementioned studies on cell line characterization and process development, it still has a limitation in predicting intracellular fluxes and identifying potential engineering targets, due to a large degree of freedom arising from an extensive and highly-interconnected metabolic network, and often exacerbated by inadequate cell culture measurements as well as a limited knowledge on intrinsic constraints and cellular objectives in complex mammalian systems. To overcome such limitation, recently, parsimonious FBA (pFBA) has been used to describe metabolic states of CHO cells (Calmels et al., 2019); the overall enzymatic fluxes are minimized to eliminate the alternative flux solutions based on the hypothesis that cells use enzymes and carbon resources efficiently, under the pressure of natural selection (Lewis et al., 2010). While this method significantly reduces flux variability compared to basic FBA, it is still unable to replicate a few key metabolic traits and physiological behaviours observed in mammalian cell culture, such as the intrinsic restriction on growth capacity and by-product secretion. More recently, another method known as “carbon constraint FBA (ccFBA)” was proposed to improve the accuracy of flux predictions in CHO cells on the basis of carbon elemental balances (Lularevic et al., 2019). Similar to pFBA, this method was designed to reduce the intracellular flux variability. However, it simply eliminates the thermodynamically infeasible pathways without any further consideration for pathway usage and enzymatic regulatory constraints which have been argued often important for biologically relevant simulation. Therefore, it is essential to incorporate the reaction kinetic information within FBA framework for elucidating the metabolic states. Indeed, the inclusion of a physical capacity limit for metabolic enzymes has been successfully shown to reduce the potential solution space in a biologically-meaningful manner (Adadi et al., 2012; Beg et al., 2007; Sánchez et al., 2017; Shlomi et al., 2011). Hence, now we can add relevant kinetic information to CHO GEM for limiting the overall metabolic enzyme usage, as such accurately identifying key bottlenecks under various cell culture conditions. By doing so, we could develop the most comprehensive and biochemically-consistent CHO GEM to-date, *i*CHO2291, from the previous model (Hefzi et al., 2016). Notably, this model includes the necessary kinetome parameters such as turnover number (*k*_*cat*_) and molecular weight of all enzymes, to enable its ready usage during the flux simulations. We then demonstrate the model utility and framework efficacy via a case study where the mechanisms underlying differential lactate overflow between two CHO clones are deciphered (Hong et al., 2018b).

## 2. Results

### 2.1 Update of CHO genome-scale metabolic model

We significantly expanded the previous CHO genome-scale model, *i*CHO1766 (Hefzi et al., 2016) following a six step procedure (**Materials and methods**). Initially, we identified the metabolite and reaction inconsistency in *i*CHO1766. Here, it should be noted that *i*CHO1766 has been built from various human GEMs including Recon2 (Thiele et al., 2013) which was previously developed based on several resources such as Recon1 (Duarte et al., 2007) and EHMN (Ma et al., 2007). Thus, some metabolites and reactions were duplicated with different identifiers across various sources. Similarly, some redundant reactions were also found in both lumped and multi-step forms. In this regard, a total of 26 duplicate metabolites and more than 250 reactions were corrected in the model, specifically for fatty acid biosynthesis, elongation and degradation pathways. Subsequently, we updated the GPR association of all reactions using the latest genome annotations, mainly focussing on two aspects: 1) formulation of proper GPR based on accurate isozyme/subunit information and 2) reaction compartmentalization. For example, if the same reaction is catalysed by two isozymes in different subcellular locations, then we connected it with the associated genes accordingly. Since the information on subcellular localization of gene products is scarce for Chinese hamster, we referred to human and mouse for corresponding CHO orthologues, thus adding more than 1400 reactions including 165 new genes. We also newly included the reactions representing the aminoacyl-tRNA charging, protein synthesis from aminoacyl-tRNAs, DNA and RNA synthesis via respective polymerases, tRNA degradation and O-glycan biosynthesis. Finally, the model expansion was completed by identifying the metabolic gaps which were filled by either adding the missing reactions from latest human GEM to close the knowledge gaps or including exchange/sink reactions to allow for the material exchange.

The resulting CHO GEM, *i*CHO2291, contains 2291 genes, 5587 reactions and 3972 metabolites. The model is available for download as Systems Biology Markup Language (SBML) file (level 3, version 1) from BioModels database (Glont et al., 2018) with identifier MODEL1912180001. A comparison of *i*CHO2961 with its predecessor (*i*CHO1766) clearly highlights a significant improvement in the number of genes and gene associated reactions, e.g., several metabolic pathways such as cholesterol metabolism, fatty acid activation, elongation and desaturation, glycerophospholipid metabolism, and N- and O-glycan biosynthesis (**Figs. 1a** and **1b**). Apart from the increased scope, we could largely reduce the numbers of blocked reactions (13% from 24%) and dead-end metabolites (9% from 23%) compared to *i*CHO1766. We also tested the ATP production rates simulated by both models in more than 25 different carbon substrates under aerobic and anaerobic conditions, and confirmed that *i*CHO2291 allowed for more reliable predictions (Data **S1**, Supporting Information).

**Figure 1.**
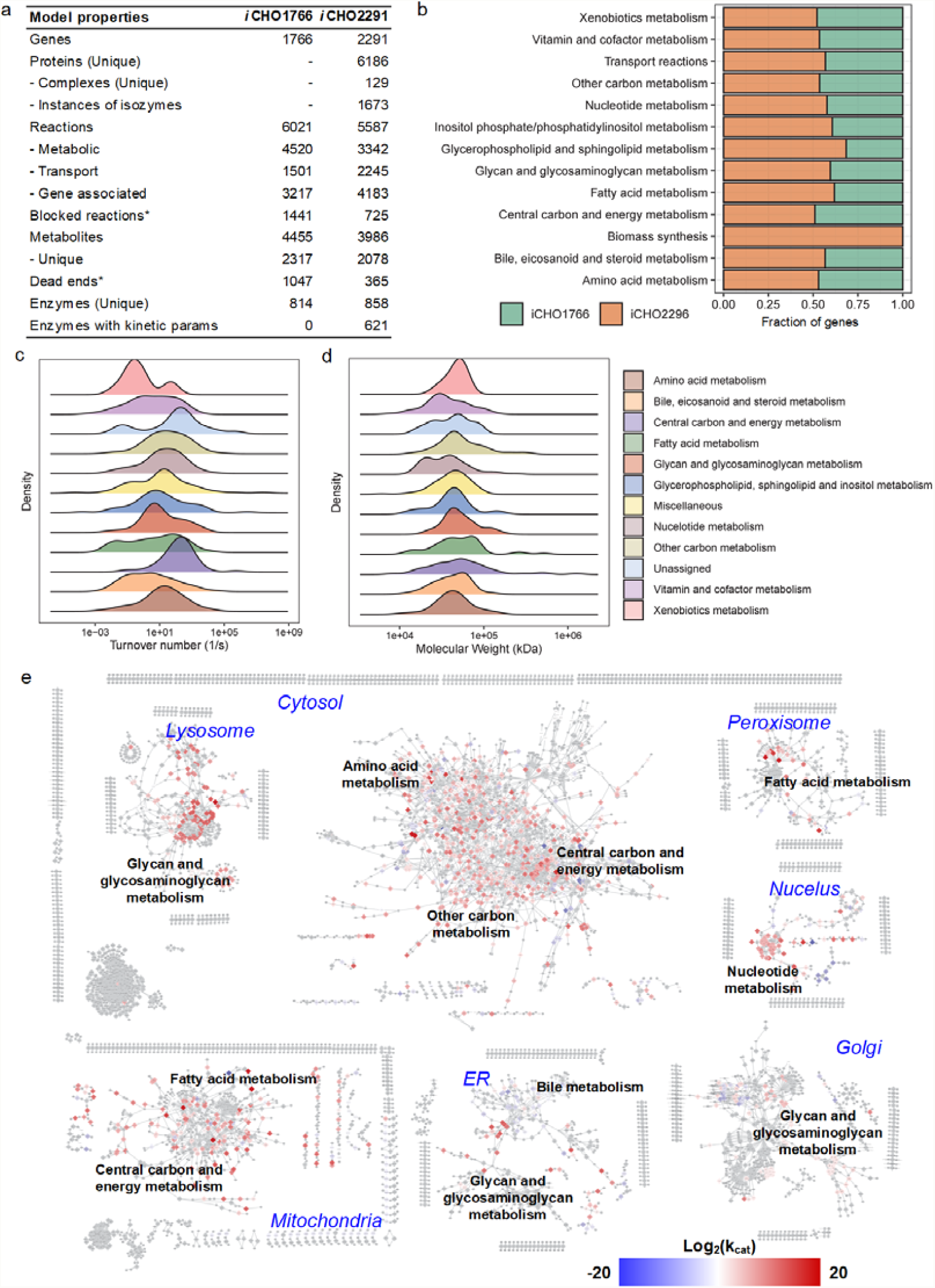
New CHO genome-scale model properties and its comparison with predecessor. a) Detailed comparison of *i*CHO1766 and *i*CHO2291 properties, b) Proportion of new genes in *i*CHO2291 when compared to *i*CHO1766, c) distribution of turnover numbers of enzymes in various pathways, d) distribution of molecular weight of various enzymes in different pathways and e) distribution of reactions with kinetic parameters visualized using the CHO genome-scale metabolic network.

Notably, *i*CHO2291 accounts for the necessary kinetome information, i.e. turnover number (*k*_*cat*_) and molecular weight, of several enzymes in Chinese hamster (see **Materials and methods;** Data **S2**, Supporting Information). To further understand the biochemical characteristics of these parameters, we explored the distribution of molecular weights and *k*_*cat*_ values. Interestingly, the *k*_*cat*_ values showed a diverse distribution with the lowest value of 1.4e-02 s^-1^ and the largest value of 2.7e+11 s^-1^. We further examined the distribution of *k*_*cat*_ values across subsystems and expectedly found that the central metabolic enzymes including glycolysis/gluconeogenesis, pentose phosphate pathway, oxidative phosphorylation and citric acid cycle have significantly higher values, indicating that these enzymes could carry out faster catalysis (**Fig. 1c**). On the other hand, *k*_*cat*_ values of N- and O-glycan synthesis and xenobiotic metabolism were markedly lower than other pathways. Here, it should be noted that since CHO cells are commonly used to produce glycosylated proteins, the N- and O-glycosylation could possibly be rate limiting. Similarly, the examination of molecular weight proteins across various subsystems showed that the biomass synthesis and central carbon and energy metabolism have bigger protein complexes than all other pathways (**Fig. 1d**).

### 2.2 Enzyme capacity constraints significantly improve intracellular flux prediction accuracy

We newly incorporated enzyme kinetic information in the updated CHO genome-scale model (*i*CHO2291) into the FBA framework in order to accurately characterize internal metabolic states of CHO cells (see **Materials and methods**). First, we compared our enzyme capacity constrained flux balance analysis (ecFBA) with previous methods, i.e. pFBA and ccFBA, in terms of the accuracy of intracellular flux predictions. To do so, we applied all three methods with heterogeneous datasets assembled from 6 different cell culture experiments where intracellular fluxes of central metabolic pathways were measured using the carbon isotope labelling. Note that we selected the datasets in such a way that we could assess both the effects of cell line engineering (Templeton et al., 2014) and changes in media (McAtee Pereira et al., 2018) on intracellular metabolic flux distribution. Note that data from Templeton et al. (2014) contain the intracellular flux measurements from two clones in which anti-apoptotic gene Bcl-2 was overexpressed at different levels and an untransfected CHO cell line as control, all of which grown in same chemically defined media. With the data from McAtee Pereira et al. (2018), we compared the intracellular flux distributions of the same CHO clone in three different chemically defined media conditions. Remarkably, the internal flux distributions predicted using ecFBA were highly consistent with the experimental measurements in all the six conditions compared to other two methods (**Figs. 2a - 2f**). We further examined the variations in intracellular flux values reaction-wise in order to understand how ecFBA could outperform the other two methods. While we do not observe any major differences in glycolytic and anaplerotic flux predictions among the three methods, considerable deviations were identified in TCA cycle and pentose phosphate pathway which generate most of the overall redox requirements (**Fig. 2g**). Interestingly, ecFBA significantly improves the flux predictions of the three irreversible reactions in TCA cycle, i.e. citrate synthase, isocitrate dehydrogenase and α-ketoglutarate dehydrogenase, that are also known to be allosterically regulated.

**Figure 2.**
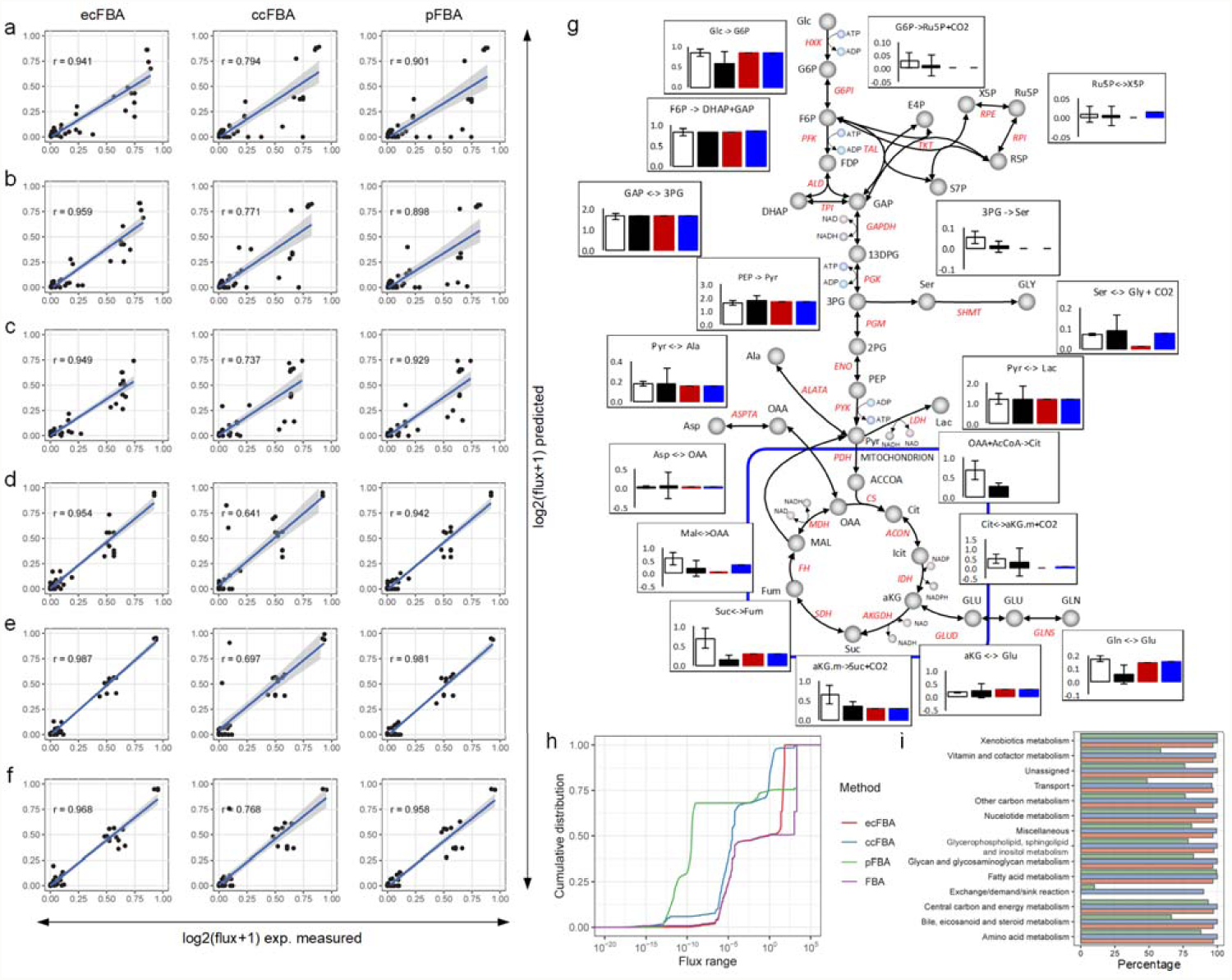
Comparison of model predictions with and without incorporation of enzyme capacity constraints and kinetic parameters. a-f) Pearson correlation coefficients of experimentally measured fluxes and the values predicted by ecFBA, ccFBA and pFBA for six datasets: (a) control condition, (b) LE clone and (c) HE clone. (d) CM media, (e) LA media and (f) LA+ media. g) Detailed comparison of experimentally predicted fluxes in (a), h) reduction of intracellular fluxes in comparison to FBA in all three methods in (a) and i) percentage reduction of fluxes in various pathways in (a). Experimentally measured fluxes in (a), (b) and (c) are from (Templeton et al., 2014) and (d), (e) and (f) are from (McAtee Pereira et al., 2018).

While all three methods have been developed to reduce the flux variability, again simulated fluxes by ecFBA showed very good agreement with experimental measurements across different datasets. Therefore, we next explored how each method performs in terms of overall flux variability reduction by comparing the flux ranges predicted in the control condition from the dataset (Templeton et al., 2014). This comparative analysis showed that ccFBA reduces the flux variability very efficiently than ecFBA and pFBA (**Fig. 2h**), although it has the lowest Pearson correlation (*r*) with experimentally observed values, thus indicating only flux variability reduction is not sufficient to predict the actual intracellular flux distribution whereas the use of actual pathways/reactions including their kinetic parameters is highly essential. We then checked whether the reduction of fluxes through any particular pathway contributes to the improved predictions in ecFBA by comparing the average flux variability reduction in each pathway. Expectedly, ccFBA resulted in maximum flux reduction across many pathways (**Fig. 2i**) while ecFBA particularly reduced the average variability much better in transport reactions, highlighting that the consumption of various nutrients is mainly limited based on the overall enzyme capacity.

### 2.3 Incorporation of enzyme capacity constraints accurately predicts the wasteful metabolism observed in CHO cell cultures

It is interesting to investigate how enzyme capacity is incorporated in ecFBA so that the “wasteful” metabolism commonly observed in CHO cell cultures can be simulated effectively. Under nutrient rich conditions, CHO cells consistently secrete toxic metabolites such as lactate and ammonia which substantially affect the cellular growth and productivity (Young, 2013). Interestingly, CHO cells use a low yielding fermentative pathway even in aerobic conditions, which generates only two molecules of ATP per mole of glucose catabolized into lactate instead of the high yielding oxidative phosphorylation providing up to 36 moles of ATP. Therefore, we examined the phenotypic behaviours of CHO cells by gradually increasing the glucose uptake from 0 to 1 mmol/gDCW/hr while simultaneously constraining the availability of all amino acids at 0.1 mmol/gDCW/hr using FBA/ccFBA/pFBA and ecFBA. It should be noted that we did not constrain oxygen availability and allowed for the free transport of all inorganic nutrients such as water, proton, sulphate and phosphate in these simulations. Not surprisingly, FBA predicted the growth rate to increase linearly with glucose uptake rates since it does not have any restriction on overall enzyme usage (**Fig. 3a**) while ecFBA simulations showed growth rate to increase only at low range of glucose uptake and then slight decrease with any further surge in glucose uptake rates. Hence, ecFBA simulations correctly predicted the experimentally observed growth pattern even without an upper limit in oxygen uptake rate, thus suggesting that additional constraints based on enzyme capacity constraints are sufficient to make realistic predictions.

**Figure 3.**
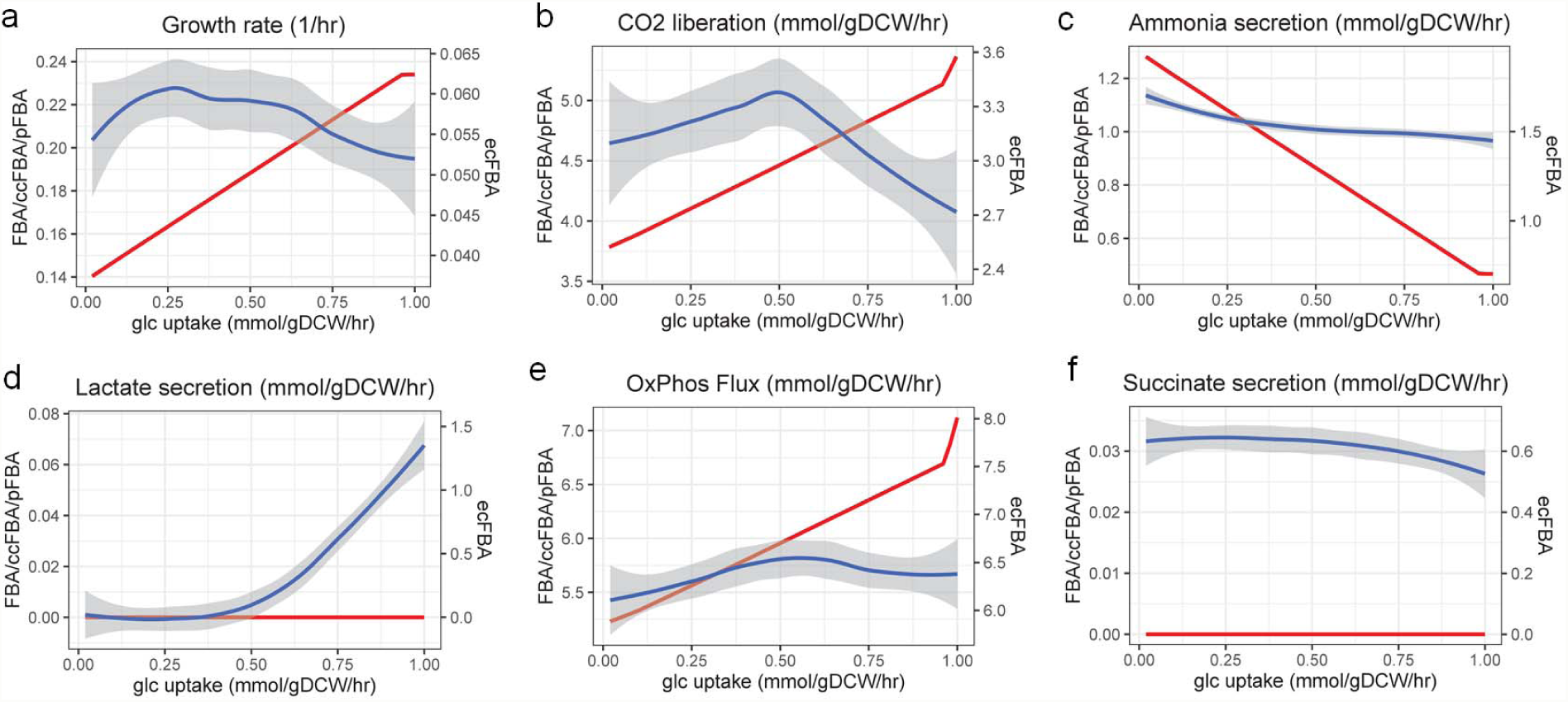
Comparison of model predictions with and without incorporation of enzyme capacity constraints and kinetic parameters over a range of glucose uptake rates. While the predicted value is plotted as it is for FBA/pFBA/ccFBA, the LOWESS smoothened trend is plotted for ecFBA. Such smoothening of simulation outcomes was performed since the original values, which are mean of 5000 simulations was noisier although it showed an overall trend.

Since FBA and ecFBA simulations predicted contrasting trends for growth rates, we further examined the secretion rates of various by-products including CO_2_, ammonia, and lactate predicted by these two methods (**Figs. 3b - 3f**). Similar to growth rate, CO_2_ secretion in FBA simulations increased linearly with increasing glucose uptake rates, indicating the possible use of oxidative phosphorylation based on its full metabolic capacity. However, ecFBA predicted the CO_2_ secretion to marginally increase until a certain point and then decreased, showing that the maximal usage of oxidative phosphorylation is possibly limited by the enzyme capacity (**Fig. 3b**). Since oxidative phosphorylation proteins have relatively high molecular weights (**Fig. 1c**), it is costlier for cells to synthesize such proteins and thus it may rely on alternative pathways which are resourceful in terms of mass even though they are not energetically efficient. Interestingly, this point at which the CO_2_ flux starts to decrease also marked the onset of lactate secretion in ecFBA simulations which could not be captured by FBA (**Fig. 3d**). Such condition clearly reflects the typical aerobic glycolysis: when glucose is available in abundant quantities, it is partially oxidized depending on the enzyme availability while the rest is metabolized into lactate (Vazquez et al., 2010). Note that any further increase in glucose gave rise to exponentially increased lactate secretion, indicating that a higher proportion of fermentative pathway is being used to produce more ATP even under aerobic condition since the amount of enzyme total mass is limited and thus the cell has to channel substantially higher flux to compensate for the lower ATP to carbon yield.

### 2.4 ecFBA delineates the clone- and media-specific lactate overflow metabolism in CHO cells

With the high confidence of *i*CHO2291 model quality and prediction accuracy by ecFBA in simulating the overflow metabolism, we analysed the lactate metabolism of two CHO clones which showed interesting trends in different media (Hong et al., 2018b). Briefly, clone A exhibited less lactate secretion in exponential phase and consumed lactate in stationary phase regardless of media, while clone B produced higher lactate and continuously secreted lactate in stationary phase in the in-house medium (m1). Interestingly, both clones consumed lactate in the stationary phase under a mixed medium containing a mixture of CD-CHO and PF-CHO at 1:1 ratio (m2). We chose the exponential phase data (day 1 to day 3) from the two clones (A and B) in media 1 and 2, i.e. Am1, Am2, Bm1 and Bm2, for further analysis (**Figs. 4a - 4f**).

**Figure 4.**
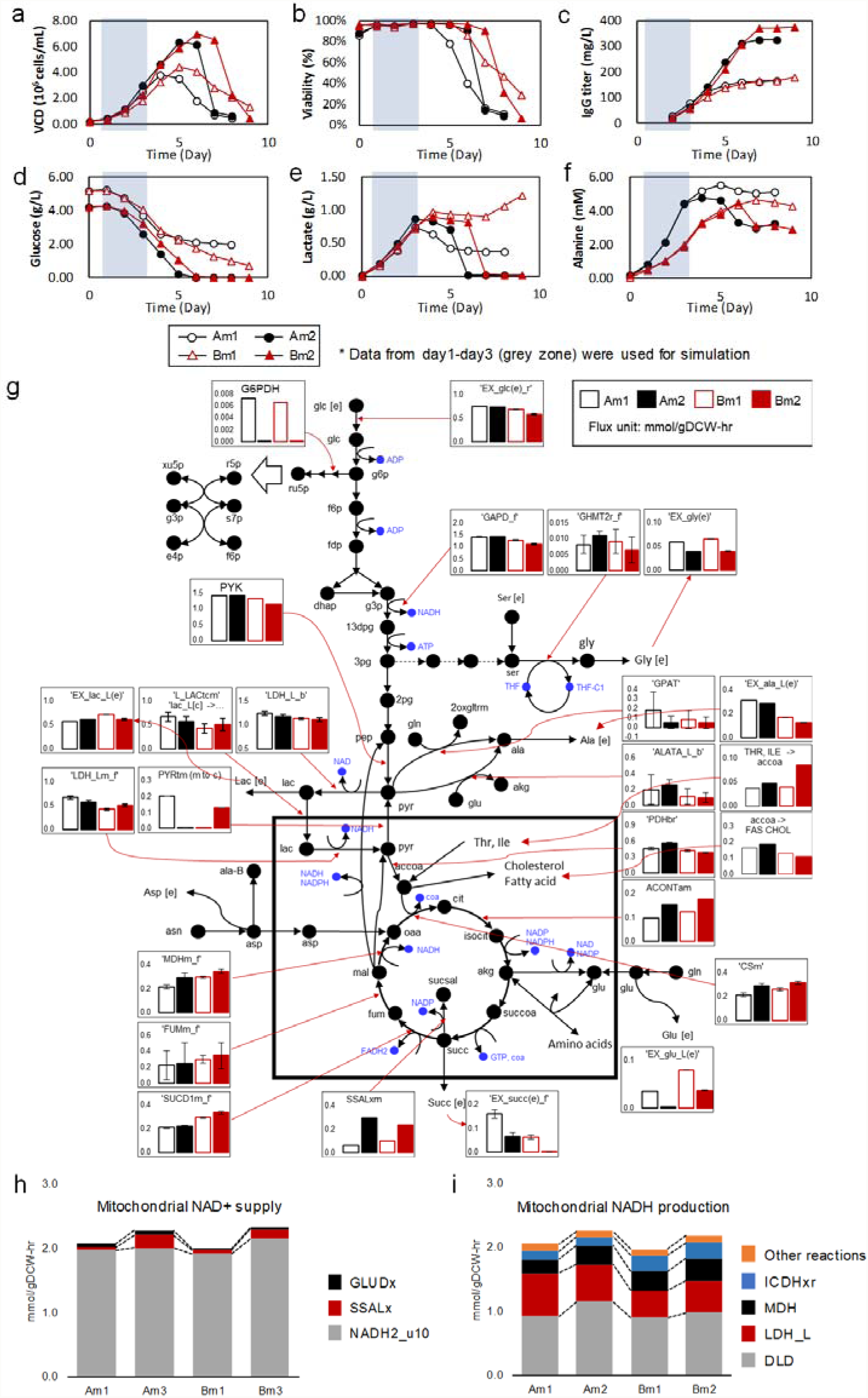
Key differences in the metabolic fates of two CHO clones in two different media. (a) Viable cell density, (b) %viability, (c) IgG titre, (c) glucose, (d) lactate, (e) alanine concentrations in batch cultures. (g) simulation results of a few key reactions in central metabolism whose flux distributions are significant are shown, (h) contribution of reaction fluxes to mitochondrial NAD supply and (i) mitochondrial NAD production.

#### 2.4.1 Clone-specific lactate metabolism

Compared to clone B, clone A showed a slightly higher glycolysis fluxes in both media. Based on the ecFBA simulations, we found that alanine was produced by two routes via: 1) alanine transaminase (ALATA) from glutamate and pyruvate and 2) glutamine pyruvate aminotransferase (GPAT) from glutamine and pyruvate. While ALATA showed comparable fluxes, GPAT flux is affected by nutrient conditions (**Fig. 4g**). Apart from alanine fluxes, ecFBA simulations captured the significantly different metabolic fates of lactate between two clones. In clone A, more cytosolic lactate (cLac) was transported into mitochondria (54% in Am1 vs 37% in Bm1), thus secreting less lactate in spite of higher net lactate generation. Subsequently, this mitochondrial lactate (mLac) was re-oxidized to mitochondrial pyruvate (mPyr) by mitochondrial lactate dehydrogenase (mLDH) (**Fig. 4g**). This lactate transport into mitochondria and reoxidation enables clone A to achieve higher energy production from the increased glycolysis, supplying more cytosolic NAD+ (cNAD+) from mitochondria. While clone A increased mLDH flux requiring mitochondrial NAD+ (mNAD+), we observed reduced fluxes through other reactions along the TCA cycle such as malate dehydrogenase (MDH) and isocitrate dehydrogenase (ICDH). This possible scenario suggests that lactate re-oxidation in mitochondria or extracellular secretion is determined by the availability of mNAD+.

#### 2.4.2 Media-specific lactate metabolism

Since medium 2 resulted in better culture performance in terms of growth and lactate overflow metabolism regardless of clonal variation, internal metabolic fluxes between m1 and m2 in clones A and B were compared to identify media specific effects. As mentioned earlier, glutamine derived alanine production via GPAT was significantly reduced in m2 in both clones as the amount of extracellular glutamine consumed is lesser compared to m1. More importantly, m2 also showed similar lactate secretion profiles between two clones. Further examination of intracellular metabolic states under m2 allowed us to find the less lactate transport to mitochondria in clone A compared to clone B (**Fig. 4g**). Additionally, clone A back-transported 37% of mPyr to cytosol in m1, but processed all mPyr by PDH in m2. Clone B, on contrary, showed negligible mPyr transport to cytosol in m1, while m2 showed 30% back-transport. Such differential transportation of pyruvate as well as lactate between cytosol and mitochondria is driven by combined factors including mNAD+ availability and PDH/CS enzyme capacity for subsequent metabolic processing, which simultaneously affect flux distribution in many other pathways such as TCA cycle. Interestingly, m2 showed higher mitochondrial NAD/NADH turnover rates in both clones, mostly by increasing fluxes through succinate semialdehyde dehydrogenase and glutamate dehydrogenase (**Fig. 4h**). Between the two clones in m2, Am2 utilized less NAD+ for mLDH due to increased fluxes of other NAD+-requiring reactions such as DLD (PDH and akgD) and MDH when compared to Bm2 (**Fig. 4i**). Therefore, we hypothesize that the lactate secretion phenotype is result of differential redox turnover rates under m2, which is in turn a combinatorial effect of cell line characteristics and redox response to the medium.

## 3. Discussion

In this study, we have extensively updated the previous CHO GEM (Hefzi et al., 2016) by correcting the biochemical inconsistency and including new genes, reactions and pathways. We also incorporated kinetome parameters of enzymes such as their turnover numbers and molecular weights which are fully utilized as additional constraints within the FBA framework, i.e., ecFBA, so that intracellular flux variations can be effectively reduced to accurately characterize cell culture behaviours and metabolic states. Thus, ecFBA was able to achieve such high concordance with experimentally measured fluxes mainly due to the usage of kinetic parameters and the limitation imposed on overall enzyme. Note that although both ecFBA and pFBA aim in identifying the most plausible intracellular flux states based on the pathway efficiency, they still differ remarkably in terms of operating procedure. pFBA assumes that cells use minimum resources to achieve the desired cellular state, and thus determines the most efficient pathway only based on reaction stoichiometry. However, ecFBA additionally considers the catalytic efficiency of enzymes by constraining relevant kinetic parameters as well as the constraints on total enzyme capacity to limit the amount of carbon which can be metabolized within the cell (Adadi et al., 2012; Beg et al., 2007; Sánchez et al., 2017; Shlomi et al., 2011). We also further showed that unlike classical FBA or pFBA, ecFBA could successfully predict the overflow metabolism under glucose excess conditions without explicit limitation on oxygen uptake rates, confirming the hypothesis that overflow metabolism could be a simple consequence of efficient enzyme allocation (Chen and Nielsen, 2019). Such simulations clearly highlight that the incorporation of overall enzyme capacity constraint and the use of enzyme kinetic parameters can serve as proxy for predicting the metabolic regulation exclusively based on the enzyme allocation.

Lactate overflow mechanism is one of the scientifically and industrially relevant research topics in CHO cell-based bioprocessing, because increased lactate secretion usually leads to the reduced production yields and inferior quality profiles (Young, 2013). Interestingly, the ecFBA simulations enabled us to find that lactate transport to mitochondria could be a key factor of cell culture performance, which is determined by the availability of mitochondrial NAD+ for mitochondrial LDH. Our ecFBA simulation results indicate lactate to possibly enter TCA cycle as an energy source in good agreement with the previous report (Brooks et al., 1999). Moreover, recent studies using C13-lactate isotope further found that many mammalian cell types and tissues could potentially use lactate as a major carbon source for TCA cycle and energy, with less contribution of glucose (Chen et al., 2016; Faubert et al., 2017; Hui et al., 2017). Here, we show that lactate transport is determined by the mitochondrial redox response which is, in turn, affected by the media used. Depending on the cell lines and media, required amount of NAD+ for mitochondrial LDH to oxidize lactate might be produced by relevant fluxes through other NADH-producing enzymes such as MDH and ICDH. As all these NAD+-requiring mitochondrial enzymes are located in outermembrane space in mitochondria (Passarella et al., 2014), competitive reactions and transport of lactate or pyruvate might be inter-related.

Apart from lactate being an energy source to facilitate TCA cycle, our simulations also indicated another possible scenario involving lactate-pyruvate cycling pathway between cytosol and mitochondria, i.e. cytosolic lactate → mitochondrial lactate → mitochondrial pyruvate → cytosolic pyruvate. Although such flux cycling may not be energetically efficient, this lactate transport enables cells to actively provide NAD+ in cytosol, as such maintaining glycolysis to simultaneously generate NADH and ATP in mitochondria. Interestingly, a recent study, which analysed the intracellular metabolism of CHO cells co-consuming glucose and lactate in late-growth phase, suggested two possible lactate metabolic routes: (1) external lactate → cytosolic lactate → cytosolic pyruvate and (2) external lactate → cytosolic lactate → mitochondrial lactate → mitochondrial pyruvate (Martínez-Monge et al., 2019), and highlighted that route 2 displays higher metabolic similarity with early growth phase in terms of cytosolic and mitochondrial NADH regeneration. In fact, fed-batch cultivation frequently shows such distinctive metabolic feature in the growth phase such that active glucose uptake and lactate consumption or marginal production are accompanied by active cell growth (Freund and Croughan, 2018; Konakovsky et al., 2016). Therefore, the lactate cycling could benefit CHO cells to achieve high glycolytic fluxes by efficiently balancing the cellular redox using the available nutrients in mitochondria, although its possible existence awaits experimental verification using C13-isotope labelling.

Although ecFBA simulations are capable of simulating the overflow metabolism and predicting the intracellular metabolism better than other methods, the accuracy and prediction capability of the model largely reply on the quality of kinetic parameters – turnover number (*k*_*cat*_) of each enzyme in the model. One key challenge is to collect reliable data due to their limited availability: *in vitro k*_*cat*_ values are available only for about 40% of the enzymes in the CHO GEM irrespective of organism from which the enzyme is isolated. While we have partly addressed the issue by randomly ranging the known values, machine learning approaches can be applied to predict the turnover numbers using enzyme biochemistry and protein structure (Heckmann et al., 2018). In another approach, multi-omics high-throughput data can be used to estimate *in vivo k*_*cat*_ values (Davidi et al., 2016).

Recently, digital transformation of the entire bioprocess has been discussed as future direction toward next generation biomanufacturing. Specifically, a huge amount of CPPs including pH, temperature, dissolved oxygen (DO), osmolality and nutrient profiles in the media as well as the critical quality attributes (CQAs) of the product (e.g., N-glycosylation patterns, charge variants and protein aggregation) and KPIs (e.g., titre and yield) can be on-line or at-line monitored and digitalized due to the advancements in soft sensor techniques. Next step is to manage the data, relate inputs (CPP) and outputs (CQA and KPI), and make real time predictions of the bioprocess outcomes for the enhanced cell culture performance via “Digital Twin” models which virtually replicate the whole process. It is envisioned that such bioprocess Digital Twins can be developed by combining advanced data analytics possibly with artificial intelligence and mechanistic model representing the cells for virtually simulating cell culture behaviours under various environmental conditions. In this regard, *i*CHO2291 with ecFBA provides a profound platform to accelerate the development of bioreactor Digital Twins where real time culture profiles can be input to be transformed into virtual predictions of metabolic changes, cell growth and recombinant product synthesis. Hence, the *in silico* model-based digital transformation allows us to effectively identify key engineering targets and design various media/feeding strategies to control the product quality, thereby reducing significant amount of costs and time in process development.

## 4. Materials and methods

### 4.1. Update procedure for CHO genome-scale reconstruction

The *i*CHO2291 model was reconstructed by updating the previously published iCHO1766 model in a 6-step procedure: 1) identification and removal of duplicate metabolites and reactions, 2) replacement of lumped reactions into detailed steps and removal of biochemically inconsistent reactions, 3) update of GPR based on latest genome annotations (as on 1 Nov 2018), 4) correction of GPR and reaction compartment assignment based on subcellular localization, 5) inclusion of new reactions based on new genome annotations, and 6) metabolic gap identification and resolution. We initiated the model expansion procedure by identifying duplicate metabolites which are represented in two different abbreviations and redundant reactions that are either present in different abbreviations or both in detailed and lumped form. We then corrected these inconsistencies by removing the lumped reactions and retained only the detailed form. If the reaction was present only in lumped form, it was replaced by corresponding detailed form. Most of such reactions are associated with fatty acid synthesis, elongation and oxidation. Next, we manually corrected the gene-protein-reaction associations for all reactions based on the new genome annotations. We also corrected the GPR and reaction compartment assignment based on the available literature evidence for the subcellular localization of corresponding gene in human and/or mouse. An extensive search on literature and KEGG database was subsequently performed to find new genes and associated reactions that were previously missing in the model. As a result, several new reactions in specific pathways such as O-glycan metabolism and NAD(P) repair metabolism were added to the network. Finally, the metabolic gaps were filled by the inclusion of either new reactions from human model or addition of new transport/sink/demand reactions depending on the literature evidence.

### 4.2. Enzyme capacity constrained flux balance analysis

In this work, enzyme capacity constrained flux balance analysis (FBA) based on the previous formulation (Shlomi et al., 2011) was used to analyze the metabolic phenotype of the CHO cells under various environmental conditions. The corresponding optimization problem can be mathematically represented as follows:

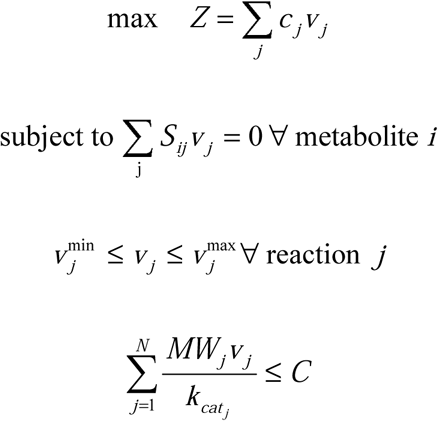

where, *Z* is the cellular objective, *c*_*j*_ is the relative weights of each metabolic reaction to biomass formation. *S*_*ij*_ is the stoichiometric coefficient of metabolite *i* of reaction *j*; *v*_*j*_ is the flux through the reaction *j*; 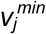 and 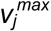 are lower and upper bounds on the flux through the reaction *j*, respectively; *MW*_*j*_ *and k*_*cat*_ are the molecular weight and turnover number of enzyme corresponding to reaction *j*. The constant C is the proportion of total enzymatic mass and is estimated based on the total protein content and the fraction of genes present in the model as described earlier (Shlomi et al., 2011).

The enzyme capacity constrained FBA was implemented using COBRA toolbox (Heirendt et al., 2019) functionalities and the biomass objective was maximized in all the simulations. In order to simulate the growth phenotype of CHO cells in different cell culture experiments where intracellular fluxes of central metabolic pathways were measured using the carbon isotope labelling, the biomass reaction was maximized while simultaneously constraining the uptake/secretion rates of glucose, lactate, ammonia and all 20 amino acids at experimentally measured values. For simulating the CHO cell growth physiology under varying glucose uptake rates, the uptake rates of all amino acids were constrained at 0.1 mmol/gDCW/hr and the glucose uptake rate was varied from 0 to 1 mmol/gDCW/hr. In both simulations, inorganic nutrients such as water, proton, sulphate and phosphate were allowed to freely flow in and out of the system. Note that when a particular reaction does not have the reactions molecular weight or *k*_*cat*_ data, then, it was assigned with random values from the permuted list of the known values. Thus, we used 5000 different permutations to obtain 5000 flux solutions for each simulation. The mean of the resulting flux solutions represents the average cellular state.

### 4.3 Compilation of parameters for enzyme capacity constraints

#### Turnover numbers

Enzyme Commission (EC) numbers were first assigned to CHO metabolic reactions based on shared reaction ID with the human Recon 2.2 model (Swainston et al., 2016). Based on the EC numbers, turnover numbers of wild-type enzymes from the BRENDA database (Chang et al., 2015) were then matched to respective reactions in the CHO model. If there was no Chinese hamster data for an enzyme, rodents (Mesocricetus spp., Phodopus spp., Rattus spp., Mus spp., and Cavia spp.), human, and others were used in the order. If there is more than one possible value, the higher turnover number is taken, so as not to over-limit the flux in the absence of more informative data.

#### Molecular weights

Using the updated gene-protein-relationship (GPR) of enzymes in the CHO GEM, we estimated their minimum molecular weight, by iterating all possible combinations of participating genes, so as not to over-restrain the flux range in the absence of discerning data. To illustrate, if the GPR stipulates that an enzyme requires gene A and B, then its molecular weight is the sum based on their translated genomic CoDing Sequences (CDS). If either gene A or B is sufficient, the selected weight is the minimum of the two translated CDS. Post-translational modifications are assumed to cause negligible weight changes. Molecular weights were directly computed based on amino acid sequences from the RefSeq database (O’Leary et al., 2016).

## Author contributions

H.C.Y., M.L. and D.-Y.L. conceived the project. H.C.Y. and M.L. reconstructed the genome-scale models, collected relevant biochemical data, developed *in silico* methods and implemented them. H.C.Y., J.H. and M.L. were involved in culture data pre-processing, modelling results interpretation and analysis. H.C.Y. and J.H. drafted the initial manuscript. M.L. and D.-Y.L. were involved in editing and revising the manuscript. M.L. and D.-Y.L. supervised and coordinated the project.

## Acknowledgements

This work was supported by Biomedical Research Council of A*STAR (Agency for Science, Technology and Research), Singapore, and the Next-Generation BioGreen 21 Program (SSAC, No. PJ01334605), Rural Development Administration, Republic of Korea. The authors also thank Kok Siong Ang for the assistance in collecting enzyme specific biochemical data and Lokanand Koduru for the useful discussions and technical support for implementing enzyme constrained flux balance analysis.

